# Insecticide resistance evolution and assisted gene flow interact to shape the evolution of plasticity

**DOI:** 10.64898/2026.04.26.720922

**Authors:** Eric G. Prileson, Bianca Campagnari, Brynne Ruotsalainen, René S. Shahmohammadloo, Cristian Zetina, Seth M. Rudman

## Abstract

Adaptive phenotypic plasticity can bolster fitness in changing environments, but the extent to which plasticity evolves rapidly, and which forces shape this evolutionary trajectory, is largely unknown. To empirically study the evolution of plasticity we first conducted a replicated field experiment in which *Drosophila melanogaster* populations adapted to insecticide exposure and a subset of these populations received high diversity assisted gene flow. We then reared individuals from each population across temperature and insecticide treatments in common garden to test the following questions: 1. Has prior selection and rapid adaptation of insecticide resistance led to evolved shifts in plasticity relative to naïve populations? 2. Does gene flow from genetically diverse populations contribute to adaptive plasticity evolution relative to gene flow-restricted low diversity populations? Both gene flow and prior evolution of resistance influenced the evolution of plasticity for multiple traits and were often maladaptive for resistant and gene flow-restricted populations, suggesting a trade-off between trait and plasticity evolution. Assisted gene flow minimized maladaptive plasticity potentially through relaxation of underlying epistatic or pleiotropic constraints. Together, these results demonstrate the dynamic interactions between trait evolution, the evolution of plasticity, and forces that shape genetic diversity with implications for conservation of threatened populations.

## Introduction

When environmental conditions shift over short time scales, they can exacerbate mismatches between organismal phenotypes and the environment. In the absence of compensatory mechanisms that reduce this mismatch, populations are likely to decline, potentially to extinction. To persist, populations can respond across generations via evolution but also more immediately within a generation through phenotypic plasticity, the capacity of a genotype to express different phenotypes in response to environmental variation (Figure 1A; West Eberhard, 2003; Pfennig, 2021). Following environmental shifts, these environmentally dependent traits could be adaptive if the plastic response moves the phenotype closer to new fitness optima (Merilä and Hendry, 2014; Seebacher et al. 2015; Diamond and Martin, 2021). On the other hand, plasticity might be maladaptive if the direction of plastic response opposes novel fitness optima or fails to minimize the phenotype-environment mismatch (Grether et al. 2005; Ghalambor et al. 2007; Brady et al. 2019). Complicating this picture is that across generations, plasticity itself can evolve rapidly with potentially strong effects to fitness (Figure 1A; Schlichting and Pigliucci, 1998; West-Eberhard, 2003; Chevin et al. 2012). Determining the pace, direction, and key drivers of the evolution of plasticity relative to trait evolution, therefore, is critical to understanding its role in population responses to environmental change, including potential rescue from extinction.

**Figure 1.**
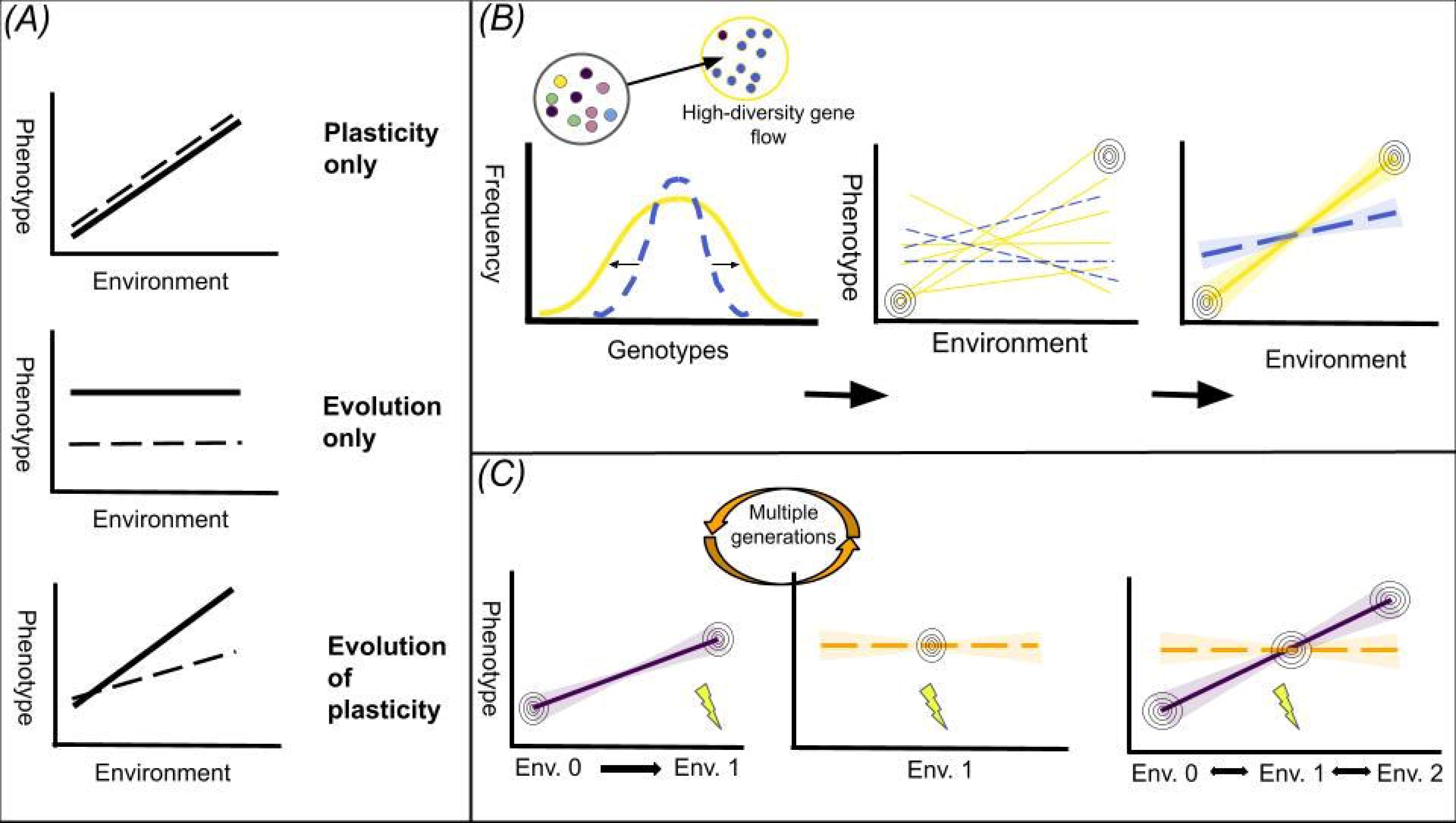
A simplified schematic of how linear reaction norms might differ between populations and processes underlying their evolution. A. Three possible outcomes for population-level reaction norms across environments. Different populations (solid and dashed lines) can both express strong plastic traits (top), evolved differences with different intercepts but with no change in slopes (middle), or different intercepts and slopes (bottom). B. The effect of genetic variation on the evolution of plasticity. An increase in the genetic diversity of a population such as through assisted gene flow (left) can provide greater variation in reaction norms in diverse (solid lines) relative to less diverse (dashed lines) populations. For genetically diverse populations (center; solid lines), there is a higher probability of reaction norms within the population matching optima across environments (concentric circles) compared to less diverse populations (dashed lines) and selection can move the average reaction norm closer to these optima (right). C. Depiction of genetic assimilation. A population can express an adaptive plastic response between an ancestral environment (Left; Env 0) and a novel one with a strong stressor (Env. 1; lightning bolt). If the novel stressor remains and the maintenance of plasticity is costly, strong selection (center) could canalize the expression of the phenotype such that the plastic response is lost over generations (orange dashed line). If the canalized trait does not match optima under returns to ancestral (right; Env 0) or new environments (Env 2), this could lead to a trade-off compared to a plastic population (purple solid line).

Plasticity can evolve if there is standing genetic variation in sensitivities to the environment (i.e., reaction norms), a process called genetic accommodation (Figure 1B; Gomulkiewicz and Kirkpatrick, 1991; Scheiner, 1993; West-Eberhard, 2003; Diamond and Martin, 2015). Fundamental evolutionary forces that alter genetic diversity then, are likely to be key for the evolution of plasticity (Barrett and Schluter, 2008; Gomulkiewicz and Houle, 2009). For example, within sub-populations, selection can decrease, whereas gene flow can increase genetic diversity on which selection can act relative to the broader metapopulation (Wright, 1930; Hartl and Clark, 1997; Gillespie, 2004). Assisted gene flow, or the directed introduction of new alleles, is one such process that can increase the diversity and probability of persistence for populations at risk of extinction (Ingvarrsson and Whitlock, 2000; Hedrick, 2004; Aitken and Whitlock, 2013; Aguilar-Gomez et al. 2025). There is conflicting evidence, however, whether increased genetic variation on its own will increase the magnitude of plasticity (Hairston et al. 2001; Sultan and Spencer, 2002; Lind et al. 2011; Hallsson and Björklund, 2012; Murren et al. 2014; David et al. 2026) or if evolved plasticity will be adaptive as gene flow could swamp locally adapted populations with maladaptive alleles (Slatkin, 1985; Lenormand, 2002; Nosil et al. 2009). In addition to potential effects of demography, the evolution of plasticity could also be constrained by costs of plasticity and maladaptive reaction norms (Snell-Rood et al. 2010; Murren et al. 2014; Ghalambor et al. 2015). Further, plasticity could also be lost altogether if selection canalizes previously plastic traits through genetic assimilation (Figure 1C; Waddington, 1953; Lande, 2009; reviewed in Scheiner and Levis, 2021). Thus, while genetic variation can lead to the evolution of plasticity, complexities of the fitness landscape will dictate the change in direction or magnitude of population-level reaction norms and their potential to rescue populations.

Natural populations face complex environments including anthropogenic stressors, but it is unclear how adaptation for one stressor bolsters or constrains the evolution of plasticity in multiple environments (Spielman et al. 2004; Pedrosa et al. 2017; Bueno et al. 2023). Spatial or temporal decoupling of a stressor in different environments presents an opportunity to test for evolutionary constraints or plastic phenotypes that are otherwise cryptic (Nyamukondiwa et al. 2011; Sinclair et al. 2013). For example, temperate insects in an agricultural environment contend with both insecticides and seasonal variation, but because insecticides are primarily applied during warm seasons, low temperatures and insecticide exposure might not co-occur. This could lead to a trade-off between insecticide resistance evolution and adaptive thermal plasticity due to negative trait correlations that oppose the direction of selection (Etterson and Shaw, 2001; Agrawal and Stinchcombe, 2009; Orr et al. 2021). In addition, exposure to both insecticide and low temperature at the same time could dramatically affect reaction norms and reveal interactions that are antagonistic or synergistic with the combined effects cancelled out or amplified, respectively (Piggot et al. 2015; Bazzicalupo et al. 2025). Given these different outcomes, there is a clear need to investigate how resistance evolution impacts the evolution of plasticity in multiple environments, but to do so requires substantial population-level replication and a system that allows for experimental manipulation.

The fruit fly *Drosophila melanogaster* (Meigen 1830) is an excellent system to test the evolution of plasticity and to identify the factors underlying it. As a model system, *D. melanogaster* is amenable to large-scale evolution experiments that provide environmental realism without reducing replication (Rudman et al. 2019; Grainger et al. 2021) and has been used to track rapid adaptation over seasonal time scales (i.e., adaptive tracking; Rudman et al. 2022).

Further, the genetic resources of *D. melanogaster* allow for manipulation of genetic variation from a known genetic history. There is also abundant empirical evidence of developmental plastic responses in thermal physiology (reviewed in Scheiner, 1993; Sørensen et al. 2016; MacClean et al. 2019) as well as genetic variation of plastic responses to low temperature (Gerken et al. 2015; Teets and Hahn, 2018) including in outdoor field settings (Mathur and Schmidt, 2017) and across latitudinal clines (Fallis et al. 2014). When reared at low temperatures, *D. melanogaster* exhibits plastic traits that are putatively adaptive including reduced fecundity (Schmidt et al. 2005; Marshall and Sinclair, 2009), improved starvation tolerance (Rion and Kawecki, 2007), greater cold tolerance (Gibert and Huey, 2001; Ayrinhac et al. 2004) and larger body sizes (Partridge et al. 1994; James et al. 1997). In addition, *D. melanogaster* can evolve resistance to insecticides (Perry et al. 2007; Karageorgi et al. 2025; Shahmohammadloo et al. 2025), but it is unknown how resistance evolution impacts the evolution of plasticity. Sudden environmental stress such as insecticide exposure can lead to genetic assimilation in *D. melanogaster* (Waddington, 1953; reviewed in Scharloo, 1991), but whether resistance is related to initial plasticity to insecticide exposure that becomes canalized is uncertain.

Here, we address how prior selection and assisted gene flow affect the evolution of plasticity in *D. melanogaster* by asking the following questions: 1. Does rapid evolution of insecticide resistance impact the evolution of plasticity relative to susceptible control populations? 2. Within insecticide-resistant populations, does gene flow from a high-diversity population facilitate the evolution of adaptive plasticity relative to populations without gene flow? To test these questions, we used populations of *D. melanogaster* that differed in prior-evolved insecticide resistance and gene flow histories. Following evolution during an outdoor growing season and common garden rearing, we raised populations across temperature, insecticide exposure, and their combination to test the evolution of plasticity in stress response and fitness-associated phenotypes. Broadly, we expected directional selection and gene flow to shift the magnitude and direction of reaction norm slopes based on changes to genetic diversity (Scheiner, 1993; Lande, 2009), but we had divergent predictions for reaction norm evolution depending on the population and developmental environment. We predicted that across temperature, constraints from selection (for resistant populations) and reduced genetic diversity (from lack of gene flow) would lead to reaction norms that were putatively maladaptive. To test if prior selection from insecticide leads to canalization, we predicted that across an insecticide treatment, resistant populations would have dampened plasticity but greater evolved trait values on the insecticide than susceptible populations. Together, these questions and our approach addressed how key processes affect the evolution of plasticity in complex environments with broader implications for responses to rapid environmental change.

## Methods

### Multigenerational experiment and founding fly populations

To investigate the evolution of plasticity, we measured plasticity in fly populations following a multigenerational outdoor selection experiment in 2024 in Vancouver, WA, USA. *D. melanogaster* populations were founded from DGRP lines (*Drosophila* Genetic Reference Panel; Mackay et al. 2012) with five males and 10 females from each line placed into a population cage. This ‘hybrid swarm’ population was allowed to mate, facilitating recombination, and grow at density controlled conditions for nine generations prior to release outdoors. This breeding design decreases linkage disequilibrium through recombination of the founder haplotypes (Weller et al. 2021). Mesocosms were maintained at the Washington State University Vancouver campus orchard (45.729 N, −122.633 W; from here on “orchard”) from June 18th until October 24th, 2024. Throughout the experiment, all populations were fed a modified Bloomington recipe media following Rudman et al. (2022).

We experimentally manipulated the genetic composition of the starting *D. melanogaster* populations. We first founded a high-diversity population from 100 DGRP laboratory fly lines (from here on ‘HD control’) and low-diversity populations from 15 DGRP lines (from here on ‘LD control’). Each of these population types were recombined in the lab and reared for four generations on control media prior to outdoor rearing. We used a subset of the LD control populations for the selection event and flies from HD control populations were used for a gene flow treatment in the outdoor seasonal experiment (see below and Figure 2).

**Figure 2.**
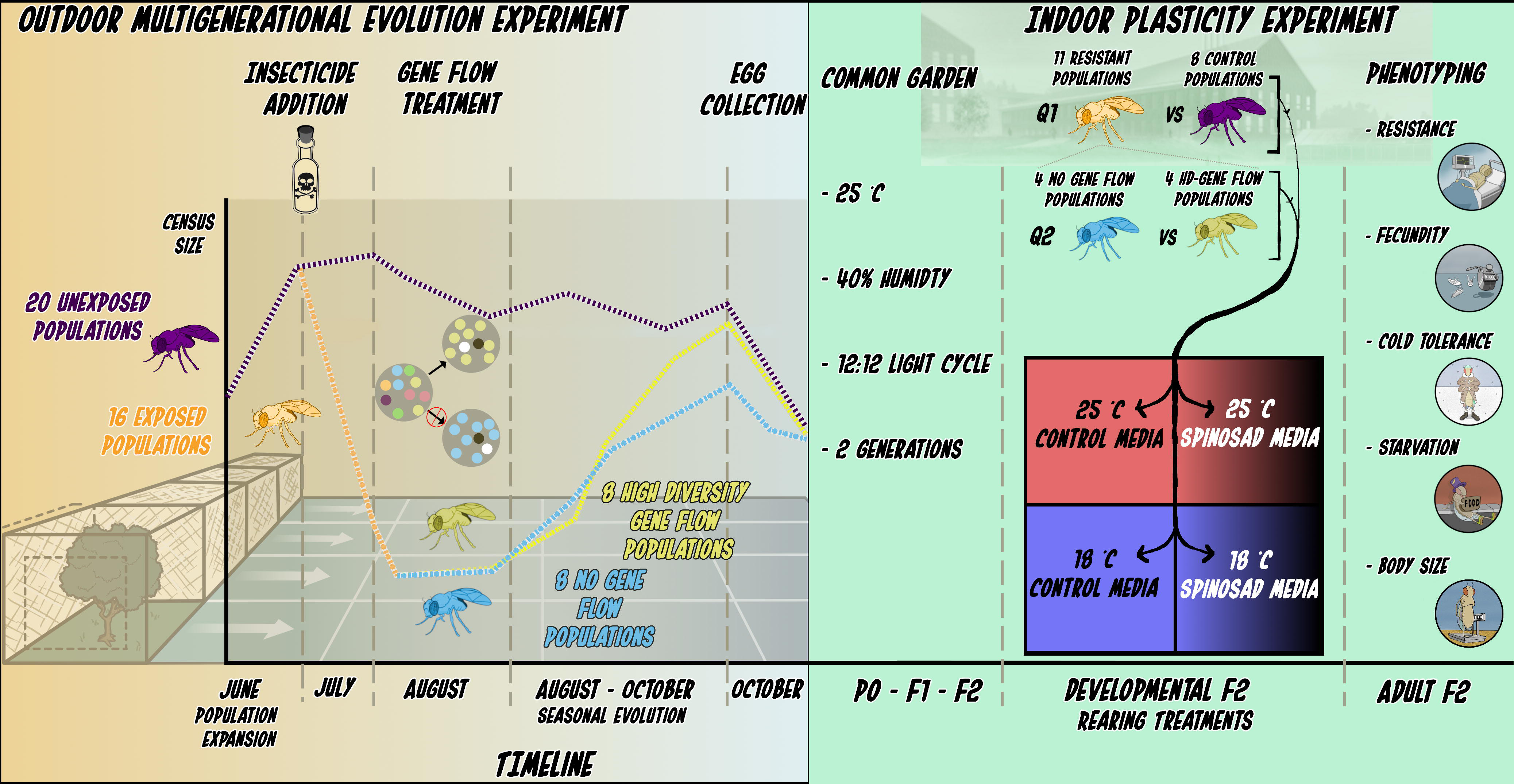
Schematic and timeline of the field study and laboratory experiment. (Left) Diagram and timeline of the multi-generational outdoor evolution experiment, showing each population type, the selection event from insecticide exposure, and gene flow treatment (see main text for details). (Right) Panels show steps of the common garden rearing, rearing treatment conditions, and phenotypes measured.

### Selection event and gene flow manipulation

We allowed populations to expand for one generation outdoors in 8 m^3^ mesocosms beginning on June 18, 2024 (2 weeks). Then, we introduced insecticide treatments to food media of 16 of the 36 populations as a strong agent of directional selection on July 5th, 2024 (Figure 2). The insecticide media was treated with a 0.0375 ug L^-1^ concentration of the organic insecticide spinosad [Entrust; Dow AgroSciences, Indianapolis, IN, USA]. This concentration corresponds with the lethal dose at which 50% of flies saw mortality based on dose-response assays conducted on founder flies under standardized laboratory conditions (see Supplementary Information in Shahmohamadloo et al. 2025). Spinosad is a widely used insecticide for fruit crops and is applied prior to and during fruit ripening (Scott et al. 2024). Spinosad targets nicotinic acetylcholine neuronal membrane receptors (nAChR), leading to neuronal overexcitation and larval mortality (Salgado, 1998; Perry et al. 2007; Martelli et al. 2022).

Following approximately one month on the insecticide treatment, during which we observed strong mortality, we initiated gene flow treatments to test the influence of gene flow on rescue (see Figure 2 and Table S1 for details on timeline). In one of the insecticide-treated groups we replaced 33% of the reproductive output from the HD control population by adding that number of eggs in a loaf pan to develop in the cage. This high diversity gene flow treatment was meant to simulate genetic rescue such that the influx of novel alleles from HD control populations increased total genetic diversity in the receiving cages. This receiving population we termed ‘HDGF’ for high diversity gene flow while a second subset of flies without gene flow we termed ‘NGF’ for no gene flow. For HDGF populations, we removed an equal proportion of eggs laid by the original population to keep the total population size consistent before and after gene flow treatment. This ensured that any change was attributable to gene flow and not direct demographic or competitive effects. NGF flies that survived exhibited population-level resistance to spinosad likely from standing genetic variation, an example of evolutionary rescue (Gomulkiewicz and Holt, 1995; Hufbauer et al. 2015). Collectively based on survival, we refer to the NGF and HDGF populations as ‘resistant’ to spinosad that we could compare to unexposed ‘control’ populations.

### Collection and initial common garden rearing

Near the end of the growing season (October 6th and 24th, 2024; Figure 2; Table S1), we collected approximately 800 eggs from each field population in density controlled 200 mL plastic bottles from 10 LD control, 10 HD control, eight NGF, and eight HDGF cages. We then reared flies indoors under common garden conditions at 25 °C under 12:12 L:D lighting and 40% humidity for three generations to control for density and environmental effects among outdoor populations. We were able to successfully rear flies from 8 LD control, four NGF and four HDGF populations. Upon eclosion of the third generation reared under common conditions, we collected 1500 eggs / 300 mL, a standard non-stressful density for the developmental plasticity experiment.

### Laboratory rearing and full factorial design

We transferred freshly laid fly eggs to media vials at a standard low density (30 eggs / vial) in one of four conditions: ([low temperature (18 °C), control media; low temperature, spinosad media; warm temperature (25 °C), control media; warm temperature, spinosad media]; Figure 2). For all combinations, we allowed all flies to develop and three to five days following eclosion, we conducted our phenotypic assays. At warm temperatures, flies developed in approximately 2 weeks and were monitored for eclosion and percent survival. Flies developing at low temperature took approximately two weeks longer and thus phenotyping occurred in two non-overlapping time periods (Table S1). Although 18 °C is a non-stressful temperature for *D. melanogaster*, it provided a lower temperature from the common garden and can induce differential gene expression and phenotypic values (Hoffman and Watson, 1993; Chen et al. 2015).

### Phenotypic assays

The following phenotypes were assayed per replicate population: 1) Insecticide resistance measured as survivorship to adulthood: the proportion of eggs (30 eggs per vial) that survived to adulthood in 3 replicate vials; 2) Fecundity: the total eggs laid by a group of five females over three days, measured daily in each of 3 replicate bottles; 3) Starvation tolerance: the time to starvation for three replicate vials containing 10 males each on agar-only media; 4) Adult body size, measured as the average dry mass of three pools of five females, dried at 55 °C for 24 hours. 5) We also included chill coma recovery time (CCRT), a static measure of cold tolerance widely used in assessing low temperature responses in *D. melanogaster* (Gibert and Huey, 2001; Rako and Hoffman, 2006). To test CCRT, we used a chill coma temperature of 4 °C for 12 hours with 15 female flies in three replicates per cage and scored recovery as the time when flies were able to right themselves at 20 °C. As flies reared at 18 °C largely did not experience chill coma at 4 °C (Table S2), we excluded this group from the cold tolerance assay and so CCRT was only compared among populations reared at 25 °C across media treatments.

### Statistical analyses

Plasticity can be modeled as the linear change in means across environmental treatments (i.e., the reaction norm slope; Via and Lande, 1985; Berrigan and Scheiner, 2004; Morrisey and Liefting, 2016) and we follow similar empirical approaches that approximate the reaction norm at the population level (Nussey et al. 2005; Charmantier et al. 2008). Therefore we constructed models with the average phenotype of each independent population as the response variable as a function of treatments. To model each phenotypic measure for each question, we used generalized linear mixed-effects models constructed with the *glmmTMB* function from the *glmmTMB* package (Brooks et al. 2017; see supplemental information for details on each model). Since the populations were common garden reared for three generations prior to developmental treatments, we consider the between-population variation in plastic responses as due to genetic differences, not within generation environmental exposures, and represent evolved differences in plasticity (Via et al. 1995; Falconer and Mackay, 1996; Berrigan and Scheiner, 2004). For each phenotypic response, a significant effect of population type, treatment condition (temperature and/or media treatment), or their interaction would indicate evolution between populations, plasticity, or the evolution of plasticity, respectively.

For our first question of resistance evolution and plasticity, we sought to test the interaction between population type, media treatment, and temperature on phenotypic response. At low temperatures and insecticide treatment, however, mortality was much higher for control populations (Table S3). This reduced sample size prevented the comparison across both media treatment and temperature for all control population phenotypes except resistance. We therefore constructed models with phenotype as the response variable, and either temperature, population, and their interaction as the predictor variables, or separately, with media treatment, population, and their interaction as predictor variables. For both types of models we considered cage (i.e., individual population) as a random intercept. To test our second question of the effect of gene flow history and plasticity, we included media treatment, population type, and temperature in our models (with the exception of CCRT). Therefore, within each model, each phenotype was the response variable and temperature, media treatment, and population type were the fixed effect predictor variables along with their pairwise interactions. As above, cage was considered a random intercept within each model.

We conducted model diagnostics using the *simulateResiduals* and *testDispersion* functions from the *DHARMa* package (Hartig and Lohse, 2022) and determined significance of the predictors using log-likelihood ratio tests in the *Anova* function with the type = “III” argument from the *car* package (Fox and Weisberg, 2019). When interaction terms were significant, post-hoc tests were conducted to identify differences between treatment levels and populations and were corrected for multiple comparisons using the *adjust = ‘fdr’* argument in the *emmeans* function from the *emmeans* package (Lenth et al. 2017). All analyses were conducted in R version 4.4.0 (R Core Team, 2025).

## Results

### Insecticide resistance and plasticity

The vast majority of flies developed successfully in three of the four treatments, but when exposed to the combination of low temperature and the insecticide, only 1.4% of flies in control populations survived to adulthood (Table S3). With this reduced sample, we could only test the evolution of plasticity in the combined low temperature and insecticide treatment for the resistance phenotype (i.e., survival when exposed). For the remaining phenotypes, we considered evolutionary divergence of plasticity across temperature and media treatments separately.

We found that plasticity of resistance evolved between resistant and control populations with a significant three way interaction between population type, temperature, and insecticide treatment (Figure 3A; estimate = 16.7 ± 4.02 SE, Χ^2^ = 17.3, P < 0.0001). This pattern was driven by considerable mortality for all populations when reared at low temperatures, but only when also developing on insecticide media (Table S4). Although survival on spinosad declined at the cooler temperature, resistant populations still maintained higher survival than control populations (Table S5).

**Figure 3.**
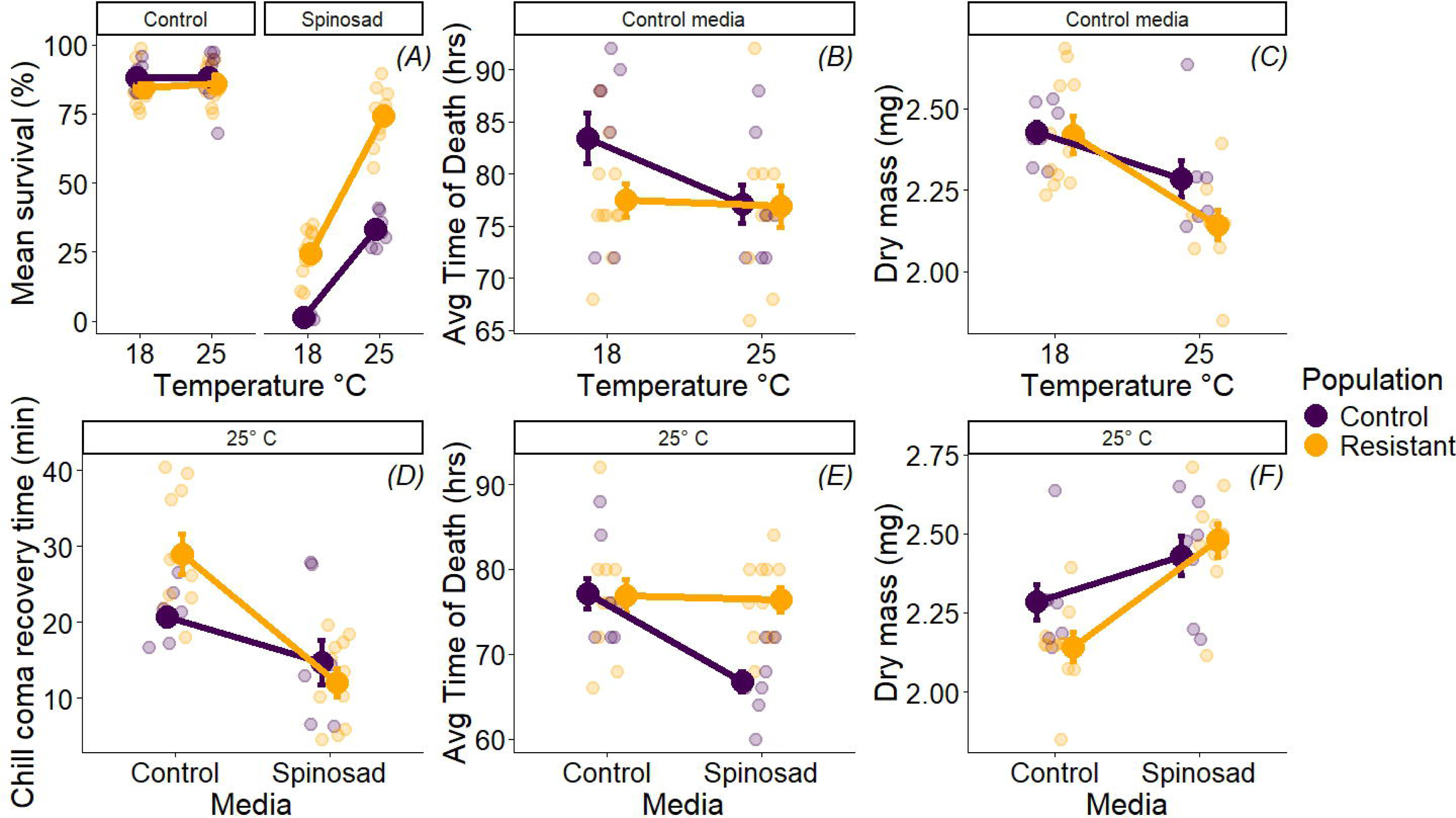
Phenotypic responses of control and resistant populations of *D. melanogaster* to temperature (top row) and media treatments (bottom row). Large points show the mean trait value of each population type ± 1 SE. Smaller points jittered behind the means show raw data values for each independent population’s phenotypic measure. A. Spinosad resistance measured as the percent egg-to-adult survival on insecticide media (note: inset panels denote media treatments). B. Starvation tolerance measured as average time to mortality. C. Average body size measured as dry mass in milligrams. D. Cold tolerance measured as chill coma recovery time in minutes. E. Starvation tolerance measured as average time to mortality across media treatment at 25 °C. F. Average body size measured as dry mass in milligrams.

#### Responses across temperature

Across temperature on control media, plasticity evolved for starvation tolerance and body size with significant interactions between population type and temperature (Figure 3B-C; Table S4,S5). Control populations exhibited stronger plasticity with improved starvation tolerance at low temperature (18 minus 25 °C = 6.35 ± 1.01 SE, t = 6.20, P < 0.0001) whereas in resistant populations low temperature did not induce a response (18 minus 25 °C = 0.559, ± 2.93 SE, t = 0.191, P = 0.851). By contrast, body size declined at a greater rate in response to warmer temperatures for resistant populations such that they exhibited a smaller size at 25 °C relative to control (control minus resistant 25 °C: estimate = 0.125 ± 0.0540 SE, t = 2.63, P = 0.0100). We did not find evidence for the evolution of plasticity for fecundity, but as expected, did see a strong plastic effect of temperature on fecundity (Figure S1; estimate = 14.0 ± 2.32 SE, X = 43.7, P < 0.0001).

#### Responses across insecticide

Across insecticide exposure at 25 °C, there was evolutionary divergence in plasticity between populations for CCRT, starvation tolerance, body size, and fecundity with a significant interaction between population type and insecticide treatment (Figure 3D-F; Table S4,S5; Figure S2). For CCRT, both populations showed faster recovery time on spinosad media (estimate - 8.58 ± 1.04 SE, Χ^2^ = 165.4, P < 0.0001), but resistant populations exhibited stronger negative plasticity with a longer recovery time on control media (Figure 3D; Control minus resistant CCRT, control media: −7.06 ± 3.44 SE, z.ratio = −2.05, P = 0.0403). Control populations exhibited strong declines in starvation tolerance when exposed to spinosad whereas resistant populations showed little plasticity (Figure 3E; control minus insecticide, control population = 10.5, ± 1.70 SE, t = 6.18, P < 0.0001; resistant = 0.478, ± 1.26 SE, t = 0.378, P = 0.708) and maintained greater starvation tolerance (Table S4). Control populations also expressed stronger declines in fecundity when exposed to insecticide (Figure S2; Control minus resistant, insecticide = −3.47 ± 1.72 SE, t = −2.02, P = 0.0439) but resistant populations exhibited a greater increase in body size from a smaller starting point on control media (Figure 3F; Resistant minus control = −0.142 ± 0.0501 SE, t = 2.83, P = 0.00550).

### Effect of gene flow on plasticity

Gene flow from the high diversity population influenced the evolution of plasticity for fecundity, starvation tolerance, and CCRT with significant interactions of population type, temperature, and/or media treatment (Figure 4A-C; Table S6). Both populations exhibited higher fecundity as temperature increased, but the induced response was weaker for NGF populations on insecticide media (Figure 4A; Table S7; HDGF minus NGF 25 °C = 5.39 ±1.54, t = −3.50, P = 0.000800). Starvation tolerance plasticity evolved on insecticide but not control media (Figure 4B; Table S6,S7) with greater starvation tolerance at 18 °C and stronger plasticity for HDGF populations across temperature compared to NGF populations (HDGF minus NGF = −10.7 ± 3.16 SE, t = −3.39, P = 0.00270). Recovery from chill coma was faster on insecticide media for both populations (Figure 4C; CCRT estimate = −17.3 ± 0.864 SE; Χ^2^ = 398.7, P < 0.0001) but on control media, HDGF populations had a faster recovery time (Table S7; HDGF minus NGF = - 10.1 ± 3.45 SE, t = 2.93, P = 0.0346).

**Figure 4.**
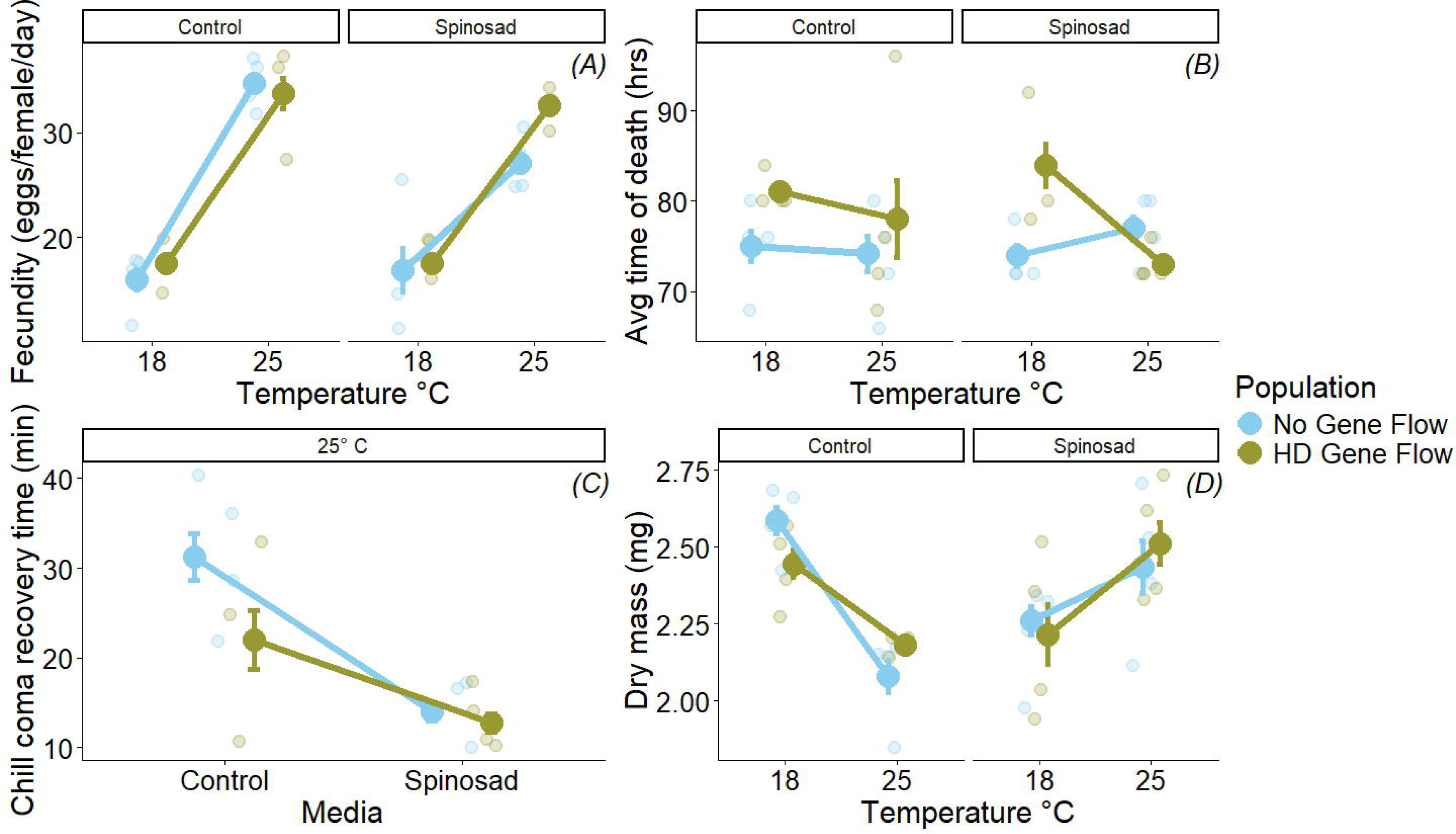
Phenotypic responses of *D. melanogaster* to temperature and media treatments (x-axis and inset panels) of resistant populations with divergent histories of gene flow: blue = NGF, gold = HDGF. Large points show the mean trait value of each population type ± 1 SE. Inset panels denote media treatments. Smaller points jittered behind the means show raw data values for each independent population’s phenotypic measure. A. Average fecundity across a three-day assay. B. Starvation tolerance measured as average time to mortality. C. Cold tolerance measured as chill coma recovery time in minutes. D. Average body size measured as dry mass in milligrams.

For body size and resistance, there were plastic responses to temperature and insecticide, but reaction norms did not differ between population types within each treatment (Figure 4D; Figure S3; Table S7). As temperature increased, body size declined for both HDGF and NGF populations (18 °C minus 25 °C no gene flow: 0.495 ± 0.0472, t = 10.5, P < 0.0001; 18 °C minus 25 °C HDGF: 0.257 ± 0.0485, t = 5.31, P < 0.0001), but on insecticide media, the direction of the plastic response was reversed (18 °C minus 25 °C no gene flow: −0.200 ± 0.0519, t = −3.86, P = 0.000200; 18 °C minus 25 °C HDGF: −0.296 ± 0.0485, t = −6.11, P < 0.0001). For resistance, both populations had high survival on control media regardless of temperature, but on insecticide, resistance declined sharply at 18 °C (Figure S4; Table S7).

## Discussion

Adaptive plasticity can be a critical component of how species respond and persist in the face of global change. Despite the potential importance of adaptive plasticity for the preservation of biodiversity, there is little known about how plasticity evolves in the complex environments typical for many natural populations. We find that plasticity does evolve rapidly. Moreover, both gene flow and selection can affect the magnitude and direction of plasticity evolution and for some phenotypes, the prior adaptation of insecticide resistance or lower diversity led to shifts that were putatively maladaptive. Together, these findings offer insight into how plasticity might evolve rapidly in fluctuating natural conditions with implications for both agriculture and conservation.

In the absence of gene flow, evolutionary rescue can proceed from standing genetic variation, but demographic costs of selection might inhibit rescue or can leave populations less capable of longer term persistence (Bell, 2013; reviewed in Carlson et al. 2014; Nordstrom et al. 2023). We found that both prior evolution of insecticide resistance and high-diversity gene flow influenced the evolution of plasticity with shifts in reaction norm slope magnitude and/or direction. Although the evolved shifts were not always in the predicted magnitude or direction, this confirmed our expectation that broad-level evolutionary processes can drive the evolution of plasticity.

Reaction norms for resistant and NGF populations in some traits were dampened and putatively maladaptive relative to control and HDGF populations respectively. The influx of alleles provided by the high-diversity gene flow treatment, however, generally alleviated maladaptive plasticity, providing empirical evidence that assisted gene flow can contribute to the evolution of adaptive plasticity and play an integral part in evolutionary rescue. We delve into the specific evolutionary shifts in reaction norms for each question below and discuss the broader implications for these results.

### Resistance evolution and the evolution of plasticity

We had predicted that prior selection from insecticides would influence the evolution of plasticity and this was supported as there were shifts in reaction norm slope magnitude and direction between control and resistant populations. Specifically, there were shifts in reaction norms for starvation tolerance and body size across temperature, as well as fecundity and cold tolerance across insecticide treatment. Since resistant populations expressed worse cold tolerance on control media and exhibited weaker starvation tolerance at the low temperature, this matched our prediction of a trade-off between resistance evolution and thermal plasticity. In addition, across insecticide treatment, resistant populations expressed dampened plasticity for fecundity and starvation tolerance, consistent with our hypothesis of canalization. Evolved shifts in reaction norms were not always maladaptive or dampened, however, as the plastic responses for body size were stronger in resistant relative to control populations and not consistent with maladaptation, suggesting that the adaptive nature of these shifts, and if plasticity is lost, is phenotype dependent.

Resistance to insecticides can evolve through many pathways and requires considering how a multifaceted genetic architecture might interact with the environment (ffrench-constant, 2013; Fournier-Level et al. 2019). Consistent with this idea, spinosad resistance in *D. melanogaster* is likely polygenic (Fournier-Level et al. 2019; Sabio et al. 2025) and reflects broad spectra of shifts in allele frequencies including (either directly or indirectly) alleles related to thermal reaction norms. For example, allelic shifts underlying resistance potentially include pleiotropy or linkage that could explain the negative correlation between resistance and cold and starvation tolerance reaction norms (Nyamukondiwa et al. 2012; Kristensen et al. 2016). Further, the intense exposure to spinosad during the growing season potentially selected for non-plastic traits in resistant populations either through canalization or because costs of maintaining plasticity was high relative to non-labile traits (Snell-Rood et al. 2010; Scheiner and Levis, 2021). Exposure to spinosad can lead to sub-lethal effects in naïve *D. melanogaster* (Weber et al. 2012; Martelli et al. 2022; Shahmohammadloo et al. 2025) and we found evidence of these effects with lower evolved fecundity and starvation tolerance for control populations.

Surprisingly, both control and resistant populations exhibited a beneficial plastic cold tolerance response when exposed to spinosad at 25 °C suggesting that development with spinosad at warm temperatures might elicit a general stress response needed for future recovery from chill coma (MacMillan et al. 2011; Sinclair et al. 2013; Cutler, 2013; Bueno et al. 2023). Considering that development at low temperature and spinosad was detrimental to survival, any effect of cross-protection of insecticide exposure seems to be limited when encountering seasonal cooling.

### Gene flow and the evolution of plasticity

As expected, gene flow played a strong role in shaping the magnitude and direction of plasticity evolution and for several traits, evolved shifts were putatively adaptive. Specifically, HDGF populations had greater fecundity at non-stressful temperatures and improved starvation tolerance at low temperatures, both consistent with adaptive life history responses (Schmidt et al. 2005; Rion and Kaweki, 2007). As in the comparison of resistant and control populations, both gene flow populations showed improved cold tolerance when exposed to spinosad and NGF populations exhibited slower, putatively maladaptive recovery time on control media.

Intriguingly, the shifts in plasticity across temperature for fecundity and starvation tolerance occurred only when populations were exposed to spinosad during development. The induced response to temperature for NGF populations was weaker (in the case of fecundity) or absent (in the case of starvation tolerance). Although any magnified effect of both treatments was alleviated in the HDGF population for these phenotypes, this was not the case for body size or resistance. Here, both populations exhibited similar plastic responses with reduced body size and survival on spinosad at low temperature, evocative of the negative synergism found in the comparison of control and resistant populations.

Why did lower diversity result in the evolution of putatively maladaptive plasticity? Given reduced standing genetic variation relative to HDGF populations, the chances of matching phenotypic optima across temperatures and media treatment were lower. This reinforces that rapid evolution of plasticity may largely be governed by similar processes as rapid evolution in other traits (Barrett and Schluter 2008), though investigations of rates of plasticity evolution, relative to rates of mean shifts in traits, could reveal more complex patterns worthy of additional study. Broadly, expanding an understanding of the relationship between genetic diversity, the rate of plasticity evolution, and the interaction between rapid trait evolution and plasticity evolution is critical to projecting organismal and population responses to environmental change.

### Implications for the evolution of plasticity in multiple environments

Insecticides can be strongly deleterious to key insect pollinators (Goulson et al. 2015; Albacete et al. 2023) and can interact with thermal physiology (Yang et al. 2018; Engell Dahl et al. 2021). In our study, when both insecticide and low temperature were experienced during development, all populations showed a strong decline in survival. This is important as effects in only one environment (e.g., insecticide exposure in warm temperature) might hide potentially negative outcomes that are only elicited when experiencing a different environment (exposure in low temperatures). Such ‘cryptic phenotypic plasticity’ has been suggested as a potential source of variation for selection to act on and indeed could impact fecundity and vital rates (Schlichting, 2008; Gibert et al. 2019). However, as we found, maladaptive effects to plasticity could also be exacerbated under differing thermal regimes and in part explain why selection for resistance might trade-off with overwintering success (Marshall et al. 2020; Prileson et al. 2025). Although this is potentially good news for managing agricultural pests, it is equally concerning for beneficial insects.

### Limitations and future directions

There are important limitations to our study that should be considered when interpreting the results. First, our design structure had only two end points of temperature and spinosad treatment (Windig et al. 2004; Chevin and Lande, 2011). This prevented us from detecting non-linear shapes of reaction norms that might differ at greater or lower environmental values (Murren et al. 2014; de Villemereuil and Chevin, 2025). Second, we did not measure direct effects to fitness; that is, we measured phenotypes across environments, but not fitness reaction norms (*sensu* Richards et al. 2006). As such, we use the language “consistent with adaptation” or “putatively (mal)adaptive” and suggest that inferences of whether reaction norm evolution was truly adaptive or not be interpreted with caution. Lastly, our population-level replication in the gene flow treatment comparison was relatively small (N = 4) and thus the mean level differences within each environment could be subject to sampling error. Future work can address each of these gaps with higher population-level replication, incorporating >2 environmental levels to measure the shape of reaction norms in continuous environments, and directly measuring fitness reaction norms through either lifetime reproduction or suitable fitness proxies.

## Conclusion

Strong selection from sudden environmental shifts can reduce genetic diversity and impact population recovery with the loss of potentially beneficial alleles, including those that are plastic (Gomulkiewicz and Holt, 1995; Bell and Gonzalez, 2008). As we show here, assisted gene flow can provide an influx of novel alleles, including those that contribute to adaptive plasticity, that temper the impact of prior selection (Scheiner and Mindell, 2020; Diamond and Martin, 2021). Indeed, this type of genetic rescue (albeit with a focus on non-plastic traits) has been at the heart of conservation efforts for threatened species such as tropical corals through assisted gene flow (Aitken and Whitlock, 2015; Baker et al. 2025). Thus, the evolution of plasticity through selection or gene flow could be a critical, but underappreciated, feature of evolutionary rescue (reviewed in Chevin et al. 2012; Tengstedt et al. 2025). The complex outcomes in multiple environments, however, underscore the need for further investigation into how plasticity evolves in a changing world.

## Acknowledgements

The authors are grateful to members of the lab for assistance with field and laboratory work and feedback on the project design. The authors also thank Simone Des Roches, Wes Dowd, Jonah Piova-Scott, Jeremiah Busch, Sarah Diamond, and Ryan Martin for helpful comments with the manuscript draft and composition.

## Data availability

Data and code scripts available on the Open Science Framework (link: https://osf.io/5yczk/overview)

