## Supplemental material and figures for "Insecticide resistance evolution and assisted gene flow interact to shape the evolution of plasticity"

**Supplementary material**

*Details on statistical analyses*

For all models, a significant effect of population type, treatment condition (temperature and/or media treatment), or their interaction would indicate evolution between populations, plasticity, or the evolution of plasticity, respectively. We constructed all models with the *glmmTMB()* function from the *glmmTMB* package (Brooks et al. 2017).

Question 1: Insecticide resistance and plasticity

Spinosad resistance:

To test the evolution of plasticity for resistance, we measured the mean survival among populations raised on spinosad and control media and at both 18 and 25 °C. For this model, we used percent survivorship as the response variable with population, temperature, and media treatment (along with their pairwise interactions) as our predictor variables, and cage as a random intercept. This model was constructed with a gaussian error distribution and the identity link function.

Starvation tolerance:

To test the evolution of plasticity for starvation tolerance across temperature, we measured the average time of survival in the absence of food among populations raised on control media and at both 18 and 25 °C. Separately, to test the evolution of plasticity for starvation tolerance across media treatment, we measured the time of survival in the absence of food among populations raised at 25 °C across both control and spinosad media. For the model across temperature, we used average hour of survival as the response variable with population, temperature, and their interaction as the predictor variables, and cage as a random intercept. Similarly, the model across media treatment also had average hour of survival as the response variable but with population, media treatment and their interaction as the predictor variables with cage as the random intercept. Both models were constructed with a gaussian error distribution and the identity link function.

Dry mass:

To test the evolution of plasticity for body size across temperature, we measured the average dry mass among populations raised on control media and at both 18 and 25 °C. Separately, to test the evolution of plasticity for body size across media treatment, we measured the average dry mass among populations raised at 25 °C across both control and spinosad media. For the model across temperature, we used average dry mass as the response variable with population, temperature, and their interaction as the predictor variables, and cage as a random intercept. Similarly, the model across media treatment also had average dry mass as the response variable but with population, media treatment and their interaction as the predictor variables with cage as the random intercept. Both models were constructed with a gaussian error distribution and the identity link function.

Fecundity:

To test the evolution of plasticity for fecundity across temperature, we measured the average eggs laid per female per day (over 3 days) among populations raised on control media and at both 18 and 25 °C. Separately, to test the evolution of plasticity for fecundity across media treatment, we measured the time of survival in the absence of food among populations raised at 25 °C across both control and spinosad media. For the model across temperature, we used average eggs laid per female per day (over 3 days) as the response variable with population, temperature, and their interaction as the predictor variables, and cage as a random intercept. Similarly, the model across media treatment also had average eggs laid per female per day (over 3 days) as the response variable but with population, media treatment and their interaction as the predictor variables with cage as the random intercept. Both models were constructed with a gaussian error distribution and the identity link function.

Chill coma recovery time:

To test the evolution of plasticity for cold tolerance across media treatment, we measured the average chill coma recovery time among populations raised at 25 °C across both control and spinosad media. For the model across media treatment also had average dry mass as the response variable but with population, media treatment and their interaction as the predictor variables with cage as the random intercept. This models were constructed with a poisson error distribution and the log-link function.

Question 2: High diversity gene flow and plasticity

Fecundity:

To test the evolution of plasticity for fecundity, we measured the average eggs laid per female per day (over three days) among populations raised on spinosad and control media and at both 18 and 25 °C. For this model, we used average eggs per female per day as the response variable with population, temperature, and media treatment (along with their pairwise interactions) as our predictor variables, and cage as a random intercept. This model was constructed with a gaussian error distribution and the identity link function.

Starvation tolerance:

To test the evolution of plasticity for starvation tolerance, we measured the average hour of mortality in the absence of food among populations raised on spinosad and control media and at both 18 and 25 °C. For this model, we used average hour of death as the response variable with population, temperature, and media treatment (along with their pairwise interactions) as our predictor variables, and cage as a random intercept. This model was constructed with a gaussian error distribution and the identity link function.

Chill coma recovery time:

To test the evolution of plasticity for cold tolerance, we measured the time to recovery among populations raised on spinosad and control media and at both 18 and 25 °C. For this model, we used recovery time as the response variable with population, temperature, and media treatment (along with their pairwise interactions) as our predictor variables, and cage as a random intercept. This model was constructed with a gaussian error distribution and the identity link function.

Body size:

To test the evolution of plasticity for body size, we measured the average dry mass among populations raised on spinosad and control media and at both 18 and 25 °C. For this model, we used average dry mass as the response variable with population, temperature, and media treatment (along with their pairwise interactions) as our predictor variables, and cage as a random intercept. This model was constructed with a gaussian error distribution and the identity link function.

Spinosad resistance:

To test the evolution of plasticity for resistance, we measured the mean survival among populations raised on spinosad and control media and at both 18 and 25 °C. For this model, we used percent survivorship as the response variable with population, temperature, and media treatment (along with their pairwise interactions) as our predictor variables, and cage as a random intercept. This model was constructed with a gaussian error distribution and the identity link function.

Figures

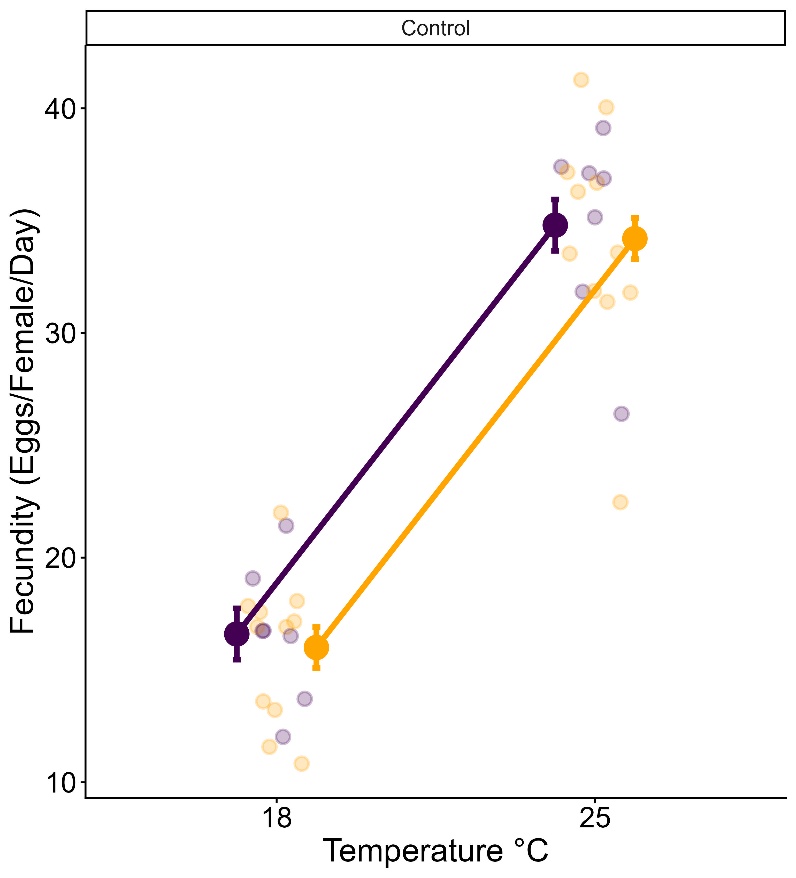

Figure S1. Fecundity (average eggs laid per female per day, for a 3 day assay) on control media measured across temperature for control and resistant populations. Large points show the mean trait value of each population type ± 1 SE. Smaller points jittered behind the means show raw data values for each independent population’s phenotypic measure.

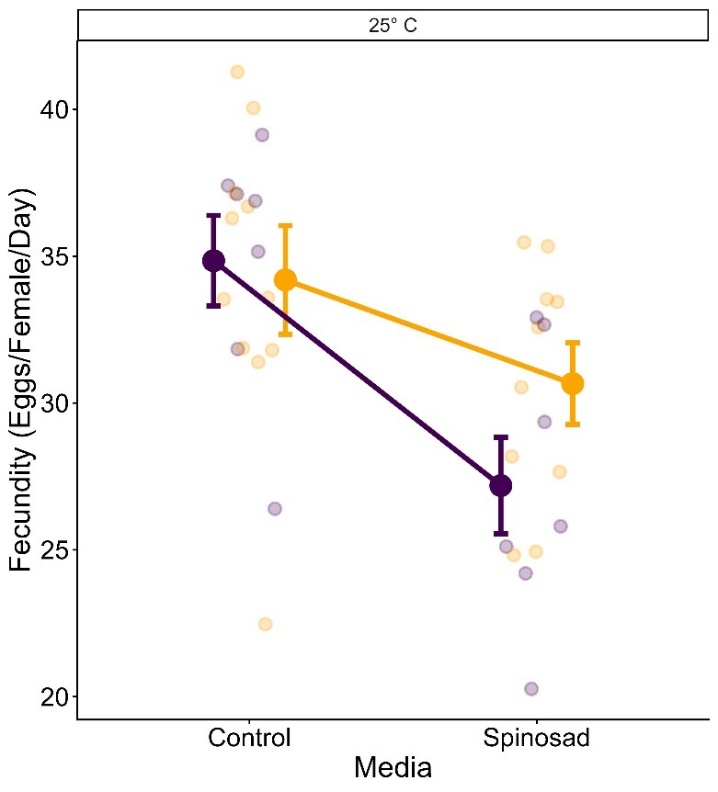

Figure S2. Fecundity (average eggs laid per female per day, for a 3 day assay) measured across media treatment at 25 °C for control and resistant populations. Large points show the mean trait value of each population type ± 1 SE. Smaller points jittered behind the means show raw data values for each independent population’s phenotypic measure.

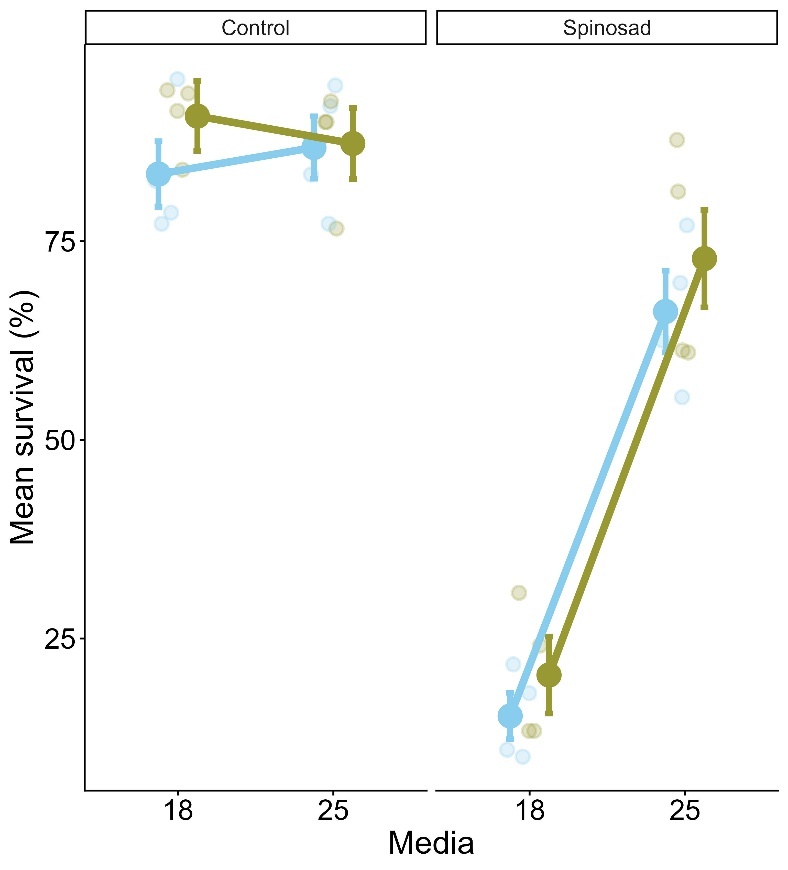

Figure S3. Spinosad resistance measured as the percent egg-to-adult survival on insecticide and control media across temperature (note: inset panels denote media treatments). Large points show the mean trait value of each population type ± 1 SE. Smaller points jittered behind the means show raw data values for each independent population’s phenotypic measure.

Tables

Table S1.

| **Date** | **Major category** | **Event** | **Steps** |
| --- | --- | --- | --- |
|  | Outdoor multigenerational experiment |  |  |
| 06/19/2024 |  | Outdoor introduction to mesocosms | Introduced all 44 populations (at this point both from 15 and 100 DGRP lines) into the orchard |
| 07/05/2024 |  | Spinosad insecticide treatments initiated | Replaced food media with spinosad treated food (0.0375 ug/L) |
| 08/15/2024 |  | Gene flow treatments initiated | Replaced original eggs laid with addition of 33% of population size with eggs from the HD control (100 DGRP outbred population) |
| 10/6/2024 |  | Fall timepoint eggs collected | 800 eggs per cage in density controlled bottles from each population type |
| 10/24/2024 |  | Secondary fall time point eggs collected | A secondary time point was needed as some of the Fall time point cages went extinct in the laboratory following collection |
|  | Common Garden |  |  |
| 10/24 - 12/22/2024 |  | Indoor common garden rearing for 3 generations (P0, F1, F2) | 2 generations of common garden rearing at 25 C, 40% humidity, and a 12:12 LD light cycle |
| 12/12/2024 |  | Transfer of eggs laid by F2 adults to the four treatments | Density controlled eggs (30 eggs / vial) to either low temperature control, low temperature spinosad, warm temperature control, and warm temperature spinosad |
|  | Round 1 Phenotyping (25 C) |  |  |
| 1/2/2025 |  | Measurement of resistance | Measured survival / successful development on each treatment |
| 1/9/2025 |  | Fecundity day 1 and starvation tolerance | First day of fecundity measures and beginning of the starvation tolerance assay |
| 1/10/2025 |  | Fecundity day 2 and CCRT | Second day of fecundity and cold tolerance assay |
| 1/11/2025 |  | Fecundity day 3 and end of starvation tolerance | Last day of fecundity measure and starvation tolerance end point |
| 1/13/2025 |  | Dry mass measure | Desiccation for 12 hours |
|  | Round 2 Phenotyping (18 C) |  |  |
| 1/15/2025 |  | Measurement of resistance | Measured survival / successful development on each treatment |
| 1/16/2025 |  | Fecundity day 1 and starvation tolerance | First day of fecundity measures and beginning of the starvation tolerance assay |
| 1/17/2025 |  | Fecundity day 2 and CCRT | Second day of fecundity and cold tolerance assay |
| 1/18/2025 |  | Fecundity day 3 and end of starvation tolerance | Last day of fecundity measure and starvation tolerance end point |
| 1/20/2025 |  | Dry mass measure | Desiccation for 12 hours |

Table S2. Proportion of flies reared at 18 C that did not undergo chill coma

| **Cage** | **Type** | **# flies in chill coma** | **Total flies** | **Percent not in chill coma** | **Treatment** |
| --- | --- | --- | --- | --- | --- |
| 2 | Control | 4 | 15 | 0.73 | Control |
| 4 | Resistant | 5 | 13 | 0.62 | Control |
| 5 | Control | 2 | 15 | 0.87 | Control |
| 6 | Resistant | 4 | 15 | 0.73 | Control |
| 8 | Control | 4 | 15 | 0.73 | Control |
| 9 | Resistant | 3 | 15 | 0.8 | Control |
| 10 | Control | 4 | 15 | 0.73 | Control |
| 11 | Resistant | 2 | 15 | 0.87 | Control |
| 11 | Resistant | 1 | 15 | 0.93 | Spinosad |
| 12 | Resistant | 4 | 15 | 0.73 | Control |
| 13 | Control | 2 | 15 | 0.87 | Control |
| 15 | Control | 5 | 15 | 0.67 | Control |
| 19 | Resistant | 2 | 15 | 0.87 | Control |
| 19 | Resistant | 0 | 12 | 0 | Spinosad |
| 20 | Resistant | 3 | 15 | 0.8 | Control |
| 22 | Control | 3 | 15 | 0.8 | Control |
| 23 | Control | 2 | 15 | 0.87 | Control |
| 27 | Control | 2 | 15 | 0.87 | Control |
| 30 | Resistant | 2 | 15 | 0.87 | Control |
| 31 | Resistant | 1 | 15 | 0.93 | Control |
| 32 | Control | 4 | 15 | 0.73 | Control |
| 33 | Resistant | 2 | 15 | 0.87 | Control |
| 34 | Control | 2 | 15 | 0.87 | Control |
| 36 | Resistant | 3 | 15 | 0.8 | Control |
| 41 | Control | 1 | 15 | 0.93 | Control |
| 42 | Control | 2 | 15 | 0.87 | Control |
| 43 | Resistant | 4 | 15 | 0.73 | Control |
| 44 | Resistant | 1 | 15 | 0.93 | Control |
| 50 | Control | 1 | 15 | 0.93 | Control |
| 54 | Resistant | 1 | 15 | 0.93 | Control |
| 60 | Resistant | 4 | 15 | 0.73 | Control |

Table S3. Proportion survival of flies within each temperature and media treatment.

| **Population type** | **Treatment** | **Temperature** | **Survival_percent** | **SE** |
| --- | --- | --- | --- | --- |
| Control | Control | 18 | 88 | 3.02 |
| Control | Spinosad | 18 | 1.39 | 0.645 |
| Resistant | Control | 18 | 84.3 | 3.36 |
| Resistant | Spinosad | 18 | 24.5 | 3.37 |
| Control | Control | 25 | 88.2 | 5.42 |
| Control | Spinosad | 25 | 33.1 | 3.11 |
| Resistant | Control | 25 | 85.8 | 2.99 |
| Resistant | Spinosad | 25 | 74.3 | 4.23 |
| HD gene flow | Control | 18 | 90.8 | 4.4 |
| HD gene flow | Spinosad | 18 | 20.4 | 4.84 |
| No gene flow | Control | 18 | 83.4 | 4.14 |
| No gene flow | Spinosad | 18 | 15.2 | 2.92 |
| HD gene flow | Control | 25 | 87.3 | 4.47 |
| HD gene flow | Spinosad | 25 | 72.8 | 6.14 |
| No gene flow | Control | 25 | 86.8 | 3.92 |
| No gene flow | Spinosad | 25 | 66.2 | 5.14 |

Table S4. Model estimates and outputs for each phenotype for the first question of insecticide resistance and the evolution of plasticity.

| **Phenotypic measure** | **Term** | **Estimate** | **SE** | **Χ^2^** | **P** |
| --- | --- | --- | --- | --- | --- |
| Resistance |  |  |  |  |  |
|  | Population | -3.68 | 3.221 | 1.31 | 0.253 |
|  | Temperature | 0.2 | 2.46 | 0.0066 | 0.935 |
|  | Treatment | -86.6 | 2.22 | 1520.3 | **< 0.0001** |
|  | Temperature : Population | 1.35 | 3.15 | 0.182 | 0.669 |
|  | Media Treatment : Pop | 26.8 | 2.84 | 89.1 | **< 0.0001** |
|  | Temperature : Treatment | 31.6 | 3.14 | 100.9 | **< 0.0001** |
|  | Population : Temperature : Media Treatment | 16.7 | 4.02 | 17.3 | **< 0.0001** |
| Starvation tolerance (Temperature; control media) |  |  |  |  |  |
|  | Population | -6.04 | 1.01 | 39.3 | **< 0.0001** |
|  | Temperature | -6.35 | 2.77 | 4.74 | **0.0295** |
|  | Population : Temperature | 5.48 | 1.29 | 18 | **< 0.0001** |
| Starvation tolerance (media treatment 25 °C) |  |  |  |  |  |
|  | Population | 0.0828 | 2.5 | 0.001 | 0.974 |
|  | Media treatment | -7.05 | 1.11 | 40.32 | **< 0.0001** |
|  | Population : Media treatment | 5.92 | 2.1 | 7.95 | **0.00481** |
| Chill coma recovery time (media treatment 25 °C) |  |  |  |  |  |
|  | Population | 7.06 | 1.14 | 3.91 | **0.0479** |
|  | Media treatment | -8.58 | 1.04 | 165.4 | **0.004517** |
|  | Population : media treatment | -7.35 | 1.04 | 74.3 | **0.00500** |
| Dry mass (Temperature; control media) |  |  |  |  |  |
|  | Population | -0.00831 | 0.0597 | 0.0133 | 0.908 |
|  | Temperature | -0.194 | 0.0715 | 22.6 | **< 0.0001** |
|  | Population : Temperature | -0.12 | 0.0912 | 12.4 | **0.000428** |
| Dry mass (media treatment; 25 °C) |  |  |  |  |  |
|  | Population | -0.138 | 0.064 | 8.03 | **0.0046** |
|  | Media treatment | 0.16 | 0.0718 | 13.4 | **0.000253** |
|  | Population : media treatment | 0.178 | 0.0909 | 14.3 | **0.000154** |
| Fecundity (Temperature; control media) |  |  |  |  |  |
|  | Population | -0.559 | 0.0403 | 0.161 | 0.688 |
|  | Temperature | 14 | 2.32 | 43.7 | **< 0.0001** |
|  | Population : Temperature | -0.131 | 0.055 | 0.0044 | 0.947 |
| Fecundity (media treatment; 25 °C) |  |  |  |  |  |
|  | Population | -0.629 | 3.09 | 0.0415 | 0.835 |
|  | Media treatment | -7.13 | 3.5 | 296.4 | **< 0.0001** |
|  | Population : media treatment | 3.54 | 4.42 | 52.6 | **< 0.0001** |

Table S5. Estimated marginal means and contrasts between levels for predictor variables for insecticide resistance and evolution of plasticity. Lower and upper confidence levels represent 95 % confidence intervals.

| **Phenotype** | **Population** | **Temperature** | **Media Treatment** | **Estimated marginal mean** | **SE** | **df** | **Lower CL** | **Upper CL** |
| --- | --- | --- | --- | --- | --- | --- | --- | --- |
| Resistance |  |  |  |  |  |  |  |  |
|  | Control | 18 | Control | 88 | 2.52 | Inf | 83 | 92.9 |
|  | Resistant | 18 | Control | 84.3 | 2.01 | Inf | 80.4 | 88.2 |
|  | Control | 25 | Control | 88.2 | 2.52 | Inf | 83.2 | 93.1 |
|  | Resistant | 25 | Control | 85.8 | 2.01 | Inf | 81.9 | 89.8 |
|  | Control | 18 | Spinosad | 1.39 | 2.28 | Inf | -3.08 | 5.86 |
|  | Resistant | 18 | Spinosad | 24.5 | 1.82 | Inf | 21 | 28.1 |
|  | Control | 25 | Spinosad | 33.1 | 2.28 | Inf | 28.7 | 37.6 |
|  | Resistant | 25 | Spinosad | 74.3 | 1.82 | Inf | 70.8 | 77.9 |
|  | **Population Contrast** | **Temperature** | **Media Treatment** | **estimate** | **SE** | **df** | **Z.ratio** | **P** |
|  | Control - Resistant | 18 | Control | 3.68 | 3.22 | Inf | 1.14 | 0.253 |
|  | Control - Resistant | 25 | Control | 2.34 | 3.22 | Inf | 0.725 | 0.468 |
|  | Control - Resistant | 18 | Spinosad | -23.1 | 2.92 | Inf | -7.93 | **< 0.0001** |
|  | Control - Resistant | 25 | Spinosad | -41.2 | 2.92 | Inf | -14.1 | **< 0.0001** |
| Starvation (temperature; control media) |  |  |  |  |  |  |  |  |
|  | **Population** | **Temperature** | **Estimated marginal mean** | **SE** | **df** | **Lower CL** | **Upper CL** |  |
|  | Control | 18 | 83.5 | 2.3 | 17.8 | 78.7 | 88.3 |  |
|  | Control | 25 | 77.1 | 2.29 | 17.6 | 72.3 | 82 |  |
|  | Resistant | 18 | 77.5 | 1.83 | 17.6 | 73.6 | 81.3 |  |
|  | Resistant | 25 | 76.6 | 1.83 | 17.8 | 72.7 | 80.4 |  |
|  | **Contrasts** | **Population** | **estimate** | **SE** | **df** | **Z.ratio** | **P** |  |
|  | Temp18 - Temp25 | Control | 6.35 | 1.02 | 17.6 | 6.2 | **< 0.0001** |  |
|  | Temp18 - Temp25 | Resistant | 0.87 | 0.812 | 17.6 | 1.07 | 0.298 |  |
| Starvation (Media treatment; 25 °C) |  |  |  |  |  |  |  |  |
|  | **Population** | **Media Treatment** | **Estimated marginal mean** | **SE** | **df** | **Lower CL** | **Upper CL** |  |
|  | Control | Control | 77.1 | 1.4 | 31 | 74.3 | 80 |  |
|  | Control | Spinosad | 66.6 | 1.55 | 41.2 | 63.5 | 69.7 |  |
|  | Resistant | Control | 76.8 | 1.13 | 32 | 74.5 | 79.1 |  |
|  | Resistant | Spinosad | 76.4 | 1.12 | 31 | 74.1 | 78.6 |  |
|  | **Contrasts** | **Population** | **estimate** | **SE** | **df** | **Z.ratio** | **P** |  |
|  | Control - Spinosad | Control | 10.5 | 1.7 | 31 | 6.18 | **< 0.0001** |  |
|  | Control - Spinosad | Resistant | 0.478 | 1.26 | 31 | 0.378 | 0.708 |  |
| Chill coma recovery time (Media treatment; 25 °C) |  |  |  |  |  |  |  |  |
|  | **Population** | **Media Treatment** | **Estimated marginal mean** | **SE** | **df** | **Lower CL** | **Upper CL** |  |
|  | Control | Control | 21.4 | 2.53 | Inf | 16.4 | 26.3 |  |
|  | Resistant | Control | 28.4 | 2.31 | Inf | 23.9 | 32.9 |  |
|  | Control | Spinosad | 14.4 | 1.78 | Inf | 10.9 | 17.9 |  |
|  | Resistant | Spinosad | 11.8 | 1.24 | Inf | 9.37 | 14.2 |  |
|  | **Contrasts** | **Treatment** | **estimate** | **SE** | **df** | **Z.ratio** | **P** |  |
|  | Control - Resistant | Control | -7.06 | 3.44 | Inf | -2.05 | 0.0403 |  |
|  | Control - Resistant | Spinosad | 2.61 | 2.16 | Inf | 1.21 | 0.227 |  |
|  | **Contrasts** | **Population** | **estimate** | **SE** | **df** | **Z.ratio** | **P** |  |
|  | Control - Spinosad | Control | 8.63 | 1.14 | Inf | 7.6 | **< 0.0001** |  |
|  | Control - Spinosad | Resistant | 16.4 | 1.45 | Inf | 11.3 | **< 0.0001** |  |
| Dry mass (temperature; control media) |  |  |  |  |  |  |  |  |
|  | **Population** | **Temperature** | **Estimated marginal mean** | **SE** | **df** | **Lower CL** | **Upper CL** |  |
|  | Control | 18 | 2.43 | 0.043 | Inf | 2.34 | 2.51 |  |
|  | Control | 25 | 2.28 | 0.042 | Inf | 2.2 | 2.37 |  |
|  | Resistant | 18 | 2.42 | 0.034 | Inf | 2.36 | 2.49 |  |
|  | Resistant | 25 | 2.14 | 0.034 | Inf | 2.08 | 2.21 |  |
|  | **Contrasts** | **Population** | **estimate** | **SE** | **df** | **Z.ratio** | **P** |  |
|  | Temp18 - Temp25 | Control | 0.143 | 0.03 | Inf | 4.75 | **< 0.0001** |  |
|  | Temp18 - Temp25 | Resistant | 0.279 | 0.024 | Inf | 11.7 | **< 0.0001** |  |
| Dry mass (media treatment; 25 °C) |  |  |  |  |  |  |  |  |
|  | **Population** | **Media Treatment** | **Estimated marginal mean** | **SE** | **df** | **Lower CL** | **Upper CL** |  |
|  | Control | Control | 2.28 | 0.039 | Inf | 2.21 | 2.36 |  |
|  | Resistant | Control | 2.14 | 0.031 | Inf | 2.08 | 2.2 |  |
|  | Control | Spinosad | 2.43 | 0.039 | Inf | 2.35 | 2.51 |  |
|  | Resistant | Spinosad | 2.48 | 0.031 | Inf | 2.42 | 2.54 |  |
|  | **Contrasts** | **Media Treatment** | **estimate** | **SE** | **df** | **Z.ratio** | **P** |  |
|  | Control - Resistant | Control | 0.142 | 0.05 | Inf | 2.83 | **0.005** |  |
|  | Control - Resistant | Spinosad | -0.05 | 0.05 | Inf | -0.998 | 0.318 |  |
| Fecundity (Temperature; control media) |  |  |  |  |  |  |  |  |
|  | **Population** | **Temperature** | **Estimated marginal mean** | **SE** | **df** | **Lower CL** | **Upper CL** |  |
|  | 18 | Control | 16.5 | 0.942 | Inf | 14.6 | 18.3 |  |
|  | 25 | Control | 34.7 | 1.37 | Inf | 32 | 37.4 |  |
|  | 18 | Resistant | 15.8 | 0.736 | Inf | 14.4 | 17.3 |  |
|  | 25 | Resistant | 34 | 1.08 | Inf | 31.9 | 36.1 |  |
|  | **Contrasts** | **Population** | **estimate** | **SE** | **df** | **Z.ratio** | **P** |  |
|  | Control - Resistant | 18 | 0.661 | 1.2 | Inf | 0.553 | 0.58 |  |
|  | Control - Resistant | 25 | 0.716 | 1.74 | Inf | 0.411 | 0.681 |  |
| Fecundity (media treatment; 25 °C) |  |  |  |  |  |  |  |  |
|  | **Population** | **Media Treatment** | **Estimated marginal mean** | **SE** | **df** | **Lower CL** | **Upper CL** |  |
|  | Control | Control | 34.8 | 1.34 | Inf | 32.2 | 37.5 |  |
|  | Resistant | Control | 34.2 | 1.07 | Inf | 32.1 | 36.3 |  |
|  | Control | Spinosad | 27.2 | 1.34 | Inf | 24.6 | 29.8 |  |
|  | Resistant | Spinosad | 30.7 | 1.07 | Inf | 28.6 | 32.8 |  |
|  | **Contrasts** | **Media Treatment** | **estimate** | **SE** | **df** | **Z.ratio** | **P** |  |
|  | Control - Resistant | Control | 0.654 | 1.72 | Inf | 0.381 | 0.703 |  |
|  | Control - Resistant | Spinosad | -3.47 | 1.72 | Inf | -2.02 | **0.043** |  |

Table S6. Model estimates and outputs for each phenotype for the second question of gene flow and the evolution of plasticity.

| **Phenotypic measure** | **Term** | **Levels** | **Estimate** | **SE** | **Χ^2^** | **P** |
| --- | --- | --- | --- | --- | --- | --- |
| Fecundity | Population |  | - | - | 1.7 | 0.427 |
|  |  | HD gene flow | 1.61 | 1.54 |  |  |
|  |  | PA gene flow | -0.0714 | 1.36 |  |  |
|  | Media Treatment | - | 1.4 | 0.735 | 3.64 | 0.0565 |
|  | Temperature | - | 18.7 | 0.653 | 821.7 | **< 0.0001** |
|  | Population : Media Treament |  | - | - | 3.05 | 0.218 |
|  |  | HD gene flow | -1.48 | 0.994 |  |  |
|  |  | PA gene flow | -0.172 | 0.886 |  |  |
|  | Population : Temperature |  | - | - | 8.82 | 0.0121 |
|  |  | HD gene flow | -2.64 | 0.924 |  |  |
|  |  | PA gene flow | -0.77 | 0.819 |  |  |
|  | Media Treatment : Temperature | - | -9.04 | 0.983 | 84.5 | **< 0.0001** |
|  | Population : Media Treatment :Temperature |  | - | - | 39.8 | **< 0.0001** |
|  |  | HD gene flow | 7.9 | 1.36 |  |  |
|  |  | PA gene flow | 6.53 | 1.21 |  |  |
| Starvation tolerance |  |  |  |  |  |  |
|  | Population |  | - | - | 5.42 | 0.0667 |
|  |  | HD gene flow | 6 | 2.63 |  |  |
|  |  | PA gene flow | 3.86 | 2.33 |  |  |
|  | Media Treatment | - | -1.3 | 2.6 | 0.248 | 0.619 |
|  | Temperature | - | -1.004 | 2.17 | 0.215 | 0.643 |
|  | Population : Media Treament |  |  |  | 1.88 | 0.392 |
|  |  | HD gene flow | 4.73 | 3.47 |  |  |
|  |  | PA gene flow | 2.29 | 3.14 |  |  |
|  | Population : Temperature |  |  |  | 0.871 | 0.647 |
|  |  | HD gene flow | -2 | 3.03 |  |  |
|  |  | PA gene flow | 0.433 | 2.69 |  |  |
|  | Media Treatment : Temperature | - | 4.3 | 3.39 | 1.61 | 0.205 |
|  | Population : Media Treatment :Temperature |  |  |  | 7.67 | **0.0216** |
|  |  | HD gene flow | -12.7 | 4.61 |  |  |
|  |  | PA gene flow | -7.58 | 4.14 |  |  |
| CCRT |  |  |  |  |  |  |
|  | Population |  | - | - | 8.84 | **0.012** |
|  |  | HD gene flow | -10.11 | 3.45 |  |  |
|  |  | PA gene flow | -3.68 | 3.03 |  |  |
|  | Treatment |  | -17.3 | 0.864 | 398.7 | **< 0.0001** |
|  | Pop*Treatment |  |  |  | 57.9 | **< 0.0001** |
|  |  | HD gene flow | 8.86 | 1.32 |  |  |
|  |  | PA gene flow | 0.536 | 1.08 |  |  |
| Dry mass | Population |  |  |  | 11.3 | **0.00356** |
|  |  | HD gene flow | -0.138 | 0.0834 |  |  |
|  |  | PA gene flow | -0.244 | 0.073 |  |  |
|  | Media Treatment | - | -0.342 | 0.0485 | 50 | **< 0.0001** |
|  | Temperature | - | -0.496 | 0.0435 | 130 | **< 0.0001** |
|  | Population : Media Treament |  |  |  | 30 | **< 0.0001** |
|  |  | HD gene flow | 0.119 | 0.0673 |  |  |
|  |  | PA gene flow | 0.306 | 0.0586 |  |  |
|  | Population : Temperature |  |  |  | 39.9 | **< 0.0001** |
|  |  | HD gene flow | 0.238 | 0.0623 |  |  |
|  |  | PA gene flow | 0.341 | 0.054 |  |  |
|  | Media Treatment : Temperature | - | 0.695 | 0.0644 | 116.4 | **< 0.0001** |
|  | Population : Media Treatment :Temperature |  |  |  | 18.6 | **< 0.0001** |
|  |  | HD gene flow | -0.142 | 0.0902 |  |  |
|  |  | PA gene flow | -0.331 | 0.0792 |  |  |
| Resistance |  |  |  |  |  |  |
|  | Population |  |  |  | 2.35 | 0.309 |
|  |  | HD gene flow | 7.3 | 5.19 |  |  |
|  |  | PA gene flow | 1.32 | 4.6 |  |  |
|  | Treatment | - | -68.2 | 3.07 | 493.7 | **< 0.0001** |
|  | Temperature | - | 3.35 | 3.41 | 0.968 | 0.325 |
|  | Pop*Treatment |  |  |  | 20.6 | **< 0.0001** |
|  |  | HD gene flow | -2.14 | 4.34 |  |  |
|  |  | PA gene flow | 13.3 | 3.85 |  |  |
|  | Pop*Temp |  |  |  | 2.02 | 0.364 |
|  |  | HD gene flow | -6.8 | 4.82 |  |  |
|  |  | PA gene flow | -2.84 | 4.27 |  |  |
|  | Treatment*Temp | - | 47.6 | 4.34 | 120.03 | **< 0.0001** |
|  | Pop*Treatment*Temp |  |  |  | 2.28 | 0.32 |
|  |  | HD gene flow | 8.3 | 6.14 |  |  |
|  |  | PA gene flow | 1.13 | 5.44 |  |  |

Table S7. Estimated marginal means and contrasts between levels for predictor variables for gene flow and the evolution of plasticity. Lower and upper confidence levels represent 95 % confidence intervals.

| **Phenotype** | **Population** | **Media Treatment** | **Temperature** | **Estimated marginal mean** | **SE** | **df** | **Lower CL** | **Upper CL** |
| --- | --- | --- | --- | --- | --- | --- | --- | --- |
| Fecundity | NGF | Control | 18 | 16 | 0.968 | Inf | 14.1 | 17.9 |
|  | HDGF | Control | 18 | 17.6 | 0.968 | Inf | 15.7 | 19.4 |
|  | NGF | Spinosad | 18 | 17.4 | 1 | Inf | 15.5 | 19.4 |
|  | HDGF | Spinosad | 18 | 17.6 | 0.975 | Inf | 15.7 | 19.6 |
|  | NGF | Control | 25 | 34.7 | 0.968 | Inf | 32.8 | 36.6 |
|  | HDGF | Control | 25 | 33.7 | 0.968 | Inf | 31.9 | 35.6 |
|  | NGF | Spinosad | 25 | 27.1 | 0.968 | Inf | 25.2 | 29 |
|  | HDGF | Spinosad | 25 | 32.6 | 0.968 | Inf | 30.7 | 34.5 |
|  | **Contrasts** |  |  |  |  |  |  |  |
|  | NGF - HDGF | Control | 18 | -1.57 | 1.37 | Inf | -1.15 | 0.251 |
|  | NGF - HDGF | Spinosad | 18 | -0.224 | 1.4 | Inf | -0.16 | 0.873 |
|  | NGF - HDGF | Control | 25 | 0.962 | 1.37 | Inf | 0.703 | 0.482 |
|  | NGF - HDGF | Spinosad | 25 | -5.47 | 1.37 | Inf | -4 | **< 0.0001** |
| Starvation tolerance | **Population** | **Media Treatment** | **Temperature** | **Estimated marginal mean** | **SE** | **df** | **Lower CL** | **Upper CL** |
|  | NGF | Control | 18 | 75 | 1.73 | 76 | 71.6 | 78.4 |
|  | HDGF | Control | 18 | 81 | 1.73 | 76 | 77.6 | 84.4 |
|  | NGF | Control | 25 | 74 | 1.78 | 76 | 70.5 | 77.6 |
|  | HDGF | Control | 25 | 78 | 1.73 | 76 | 74.6 | 81.4 |
|  | NGF | Spinosad | 18 | 73.7 | 2.26 | 76 | 69.2 | 78.2 |
|  | HDGF | Spinosad | 18 | 84.4 | 1.93 | 76 | 80.5 | 88.2 |
|  | NGF | Spinosad | 25 | 77 | 1.73 | 76 | 73.6 | 80.4 |
|  | HDGF | Spinosad | 25 | 73 | 1.73 | 76 | 69.6 | 76.4 |
|  | **Contrasts** | **Media Treatment** | **Temperature** | **estimate** | **SE** | **df** | **Z.ratio** | **P** |
|  | NGF - HDGF | Control | 18 | -6 | 2.44 | 76 | -2.46 | **0.016** |
|  | NGF - HDGF | Control | 25 | -3.99 | 2.48 | 76 | -1.61 | 0.112 |
|  | NGF - HDGF | Spinosad | 18 | -10.7 | 2.98 | 76 | -3.58 | **0.001** |
|  | NGF - HDGF | Spinosad | 25 | 4 | 2.44 | 76 | 1.64 | 0.105 |
| Chill coma recovery time | **Population** | **Media Treatment** | **Estimated marginal mean** | **SE** | **df** | **Lower CL** | **Upper CL** |  |
|  | NGF | Control | 31.3 | 2.41 | 12.7 | 26 | 36.5 |  |
|  | HDGF | Control | 21.1 | 2.46 | 13.8 | 15.8 | 26.4 |  |
|  | No_gene_flow | Spinosad | 14 | 2.41 | 12.7 | 8.77 | 19.2 |  |
|  | HD_gene_flow | Spinosad | 12.8 | 2.41 | 12.7 | 7.52 | 18 |  |
|  | **Contrasts** | **Media Treatment** | **estimate** | **SE** | **df** | **Z.ratio** | **P** |  |
|  | NGF - HDGF | Control | 10.1 | 3.45 | 13.3 | 2.93 | **0.035** |  |
|  | NGF - HDGF | Control | 3.68 | 3.03 | 12.7 | 1.22 | 0.246 |  |
|  | NGF - HDGF | Control | -6.43 | 3.07 | 13.4 | -2.1 | 0.083 |  |
|  | NGF - HDGF | Spinosad | 1.25 | 3.42 | 12.7 | 0.366 | 0.72 |  |
|  | NGF - HDGF | Spinosad | 3.14 | 3.03 | 12.7 | 1.04 | 0.72 |  |
|  | NGF - HDGF | Spinosad | 1.89 | 3.03 | 12.7 | 0.625 | 0.72 |  |
| Dry mass | **Media Treatment** | **Population** | **Temperature** | **Estimated marginal mean** | **SE** | **df** | **Lower CL** | **Upper CL** |
|  | Control | NGF | 18 | 2.58 | 0.072 | 8.6 | 2.41 | 2.74 |
|  | Spinosad | NGF | 18 | 2.23 | 0.075 | 10.1 | 2.07 | 2.4 |
|  | Control | HDGF | 18 | 2.44 | 0.073 | 8.99 | 2.27 | 2.6 |
|  | Spinosad | HDGF | 18 | 2.22 | 0.073 | 8.99 | 2.05 | 2.38 |
|  | Control | NGF | 25 | 2.08 | 0.072 | 8.27 | 1.92 | 2.25 |
|  | Spinosad | NGF | 25 | 2.43 | 0.072 | 8.27 | 2.27 | 2.6 |
|  | Control | HDGF | 25 | 2.18 | 0.072 | 8.27 | 2.02 | 2.35 |
|  | Spinosad | HDGF | 25 | 2.51 | 0.072 | 8.27 | 2.35 | 2.68 |
|  | **Contrasts** | **Population** | **Temperature** | **estimate** | **SE** | **df** | **Z.ratio** | **P** |
|  | Control - Spinosad | NGF | 18 | 0.343 | 0.053 | 73.1 | 6.52 | **< 0.0001** |
|  | Control - Spinosad | HDGF | 18 | 0.223 | 0.051 | 73.1 | 4.4 | **< 0.0001** |
|  | Control - Spinosad | NGF | 25 | -0.353 | 0.046 | 73 | -7.65 | **< 0.0001** |
|  | Control - Spinosad | HDGF | 25 | -0.33 | 0.046 | 73 | -7.16 | **< 0.0001** |
| Resistance | **Temperature** | **Population** | **Media Treatment** | **Estimated marginal mean** | **SE** | **df** | **Lower CL** | **Upper CL** |
|  | 18 | NGF | Control | 83.5 | 3.67 | 27.9 | 75.9 | 91 |
|  | 25 | NGF | Control | 86.8 | 3.67 | 27.9 | 79.3 | 94.3 |
|  | 18 | HDGF | Control | 90.7 | 3.67 | 27.9 | 83.2 | 98.3 |
|  | 25 | HDGF | Control | 87.3 | 3.67 | 27.9 | 79.8 | 94.8 |
|  | 18 | NGF | Spinosad | 15.2 | 3.36 | 19.8 | 8.23 | 22.3 |
|  | 25 | NGF | Spinosad | 66.2 | 3.36 | 19.8 | 59.1 | 73.2 |
|  | 18 | HDGF | Spinosad | 20.4 | 3.36 | 19.8 | 13.4 | 27.4 |
|  | 25 | HDGF | Spinosad | 72.8 | 3.36 | 19.8 | 65.8 | 79.8 |
|  | **Contrasts** | **Population** | **Media Treatment** | **estimate** | **SE** | **df** | **Z.ratio** | **P** |
|  | Temp18 - Temp25 | NGF | Control | -3.35 | 3.41 | 27.9 | -0.984 | 0.334 |
|  | Temp18 - Temp25 | HDGF | Control | 3.45 | 3.41 | 27.9 | 1.01 | 0.32 |
|  | Temp18 - Temp25 | NGF | Spinosad | -50.9 | 2.69 | 19.8 | -18.9 | **< 0.0001** |
|  | Temp18 - Temp25 | HDGF | Spinosad | -52.4 | 2.69 | 19.8 | -19.5 | **< 0.0001** |
